# Cyanogenic clovers invest more in rhizobia but do not affect rhizobium evolution

**DOI:** 10.64898/2026.09.11.751034

**Authors:** Julia N. Eckberg, Jennifer A. Lau, Kenneth M. Olsen, Mia M. Howard

## Abstract

Nitrogen governs the interactions between leguminous plants and their nitrogen-fixing resource mutualists, rhizobia. While most work has focused on how soil nitrogen availability shapes the ecology and evolution of the legume-rhizobium mutualism, plants’ nitrogen demands may also influence the mutualism. Plants differ in nitrogen demand, in part because of variation in specialized metabolite production, particularly in species that produce nitrogen-based defense compounds such as cyanogenic glycosides. Plants with high nitrogen demand might be expected to be more reliant on rhizobia, potentially investing more in rhizobia and exerting stronger preferences for high-quality mutualists. Here, we use white clover (*Trifolium repens*), which is polymorphic for cyanogenesis, from two biparental F_3_ mapping populations to examine whether nitrogen-based chemical defenses affect plant investment in the mutualism or rhizobium evolution. By inoculating cyanogenic and acyanogenic clover lines with a single strain of *Rhizobium leguminosarum*, we found that cyanogenic clovers produced larger root nodules than acyanogenic clover, suggesting that cyanogensis is associated with increased investment in rhizobia. Greater investment in rhizobia could select for more cooperative mutualists if accompanied by partner choice or sanctions allowing plants to preferentially reward more cooperative rhizobia. We conditioned soils with cyanogenic and acyanogenic clover lines for five plant growth cycles and compared the partner quality of rhizobia isolated from these soils. Rhizobia evolved in association with cyanogenic versus acyanogenic clover did not differ in quality. Overall, our results show that N-based anti-herbivore defense traits can influence plant investment in resource mutualists but suggest that differences in investment do not necessarily translate into long-term evolutionary changes in mutualist quality.

## Introduction

Nutritional mutualisms are shaped by the availability of traded resources. Theory and empirical work suggest that hosts persisting in high resource environments invest less in the mutualism and that over longer time scales, high resource availability should favor the evolution of hosts that are less reliant on microbial mutualists and less cooperative microbial partners (Kiers et al. 2010; Akçay & Simms 2011; Bever 2015). For example, in the legume-rhizobium symbiosis, rhizobia colonize the roots of leguminous plants through the formation of nodules and convert atmospheric nitrogen (N) into a form of bioavailable N in exchange for photosynthate. Because of the substantial carbon cost incurred by plants in this interaction, resource mutualism theory predicts that plants will only invest in symbioses with rhizobia when N is limited; consistent with this prediction, plants often form fewer nodules when fertilized with N (Streeter & Wong 1988; Luciński et al. 2002; Glyan’ko et al. 2009; Friel & Friesen 2019). Nitrogen availability can also influence the evolution of rhizobium quality. For example, rhizobia from grassland communities fertilized with N for 22 years were lower quality mutualists that provided fewer benefits to host plants compared to rhizobia isolated from unfertilized soils (Weese et al. 2015 but see Simonsen et al. 2015; Wendlandt et al. 2022).

Yet soil N-availability is likely only half of the equation; the N demands of the host could also influence both plant allocation to rhizobia and the evolution of rhizobium partner quality. Many factors influence plants’ N demands, including life history strategy (Li et al. 2024), life stage (Montesinos-Navarro 2023), and plant defense strategy—particularly in species that produce nitrogenous defense chemicals (Gleadow & Møller 2014). For example, some plants synthesize cyanogenic glycosides in their leaves, which can be broken down into reactive hydrogen cyanide molecules upon tissue damage, harming herbivores and deterring them from further damaging and consuming plant tissues (Hruska 1988; Boter & Diaz 2023). These N-based compounds are costly to produce, and as such cyanogenic plants may incur greater fitness costs when associating with low-quality partners that provide limited N benefits. Increased plant host N-demand and its potential effects on the associated host benefits of high-quality rhizobia/costs of low-quality rhizobia may influence the strength of the two primary factors that promote the maintenance of high-quality mutualists: 1) partner-fidelity fitness feedbacks and 2) partner choice or sanctions (Sachs et al. 2004). First, partner-fidelity fitness feedbacks occur when the fitnesses of the two partners are linked; more cooperative rhizobia increase host growth and fitness, so that more resources feedback to increase the fitness of the highly cooperative rhizobia. In contrast, plants associating with low-quality rhizobia are less productive and have fewer resources to allocate to their rhizobium symbionts, and both plant and rhizobium fitness are reduced (Sachs et al. 2004; Weyl et al. 2010). Because of the high N demand of cyanogenic plants, fitness feedbacks might be stronger for cyanogenic plant-rhizobium interactions than for acyanogenic plant-rhizobium interactions. Essentially, only exceptionally high-quality rhizobia may be able to support cyanogenic glucoside production and growth, whereas associations with lower quality rhizobia may not provide sufficient N to sustain both. As such, we might predict greater asymmetry in reward available to high-vs. low-quality rhizobia. In contrast, acyanogenic plants may require less N to realize growth benefits and even lower quality rhizobia may provide the growth benefits to their plant hosts that allow for carbon resources to feedback to the rhizobia.

Similarly, the strength of partner choice and sanctions may also be affected by N demand. Partner choice occurs when the host selectively forms symbioses with rhizobia that provide the greatest resource/fitness benefits to the host, while sanctions occur when hosts provide fewer resources to lower quality rhizobia postinfection (Sachs et al. 2004). Partner choice and sanctioning are genetically variable (Simonsen & Stinchcombe 2014), which may be in part due to genetic variation in N demand. For example, we might expect that cyanogenic plants would more rigorously discriminate partner quality when forming symbioses and preferentially associate with more beneficial partners or impose more severe sanctions on uncooperative partners.

Here, we use white clover (*Trifolium repens*), a plant that is polymorphic for cyanogenesis, to investigate how N-based chemical defenses affect plant investment in N-fixing rhizobium mutualists at the organismal and evolutionary scale. Given that the concentration of N-based defense compounds such as cyanogenic glucosides in plant tissues is highly sensitive to N availability (Gleadow & Møller 2014; Imakumbili et al. 2019), the capacity of plants to produce nutritionally expensive chemical defenses may shape investment in or quality of rhizobia. To test whether cyanogenic plants invest more in resource mutualisms, first we inoculated cyanogenic and acyanogenic genotypes of clover with *Rhizobium leguminosarum* and compared investment in rhizobium symbionts. Then, we conducted an experimental evolution study where we evolved replicated rhizobium populations in the presence of cyanogenic or acyanogenic clover genotypes for five plant growth cycles before isolating strains of *R. leguminosarum* and comparing their quality as plant mutualists. Specifically, we ask two interrelated questions:

1. Does cyanogenesis affect clover investment in rhizobium mutualists?
2. Does cyanogenesis in clover affect the evolution of rhizobium quality?

## Methods

### Experiment 1: Investment in rhizobia across plant cyanotype

#### Experimental setup

To investigate differences in rhizobium investment between clover cyanotypes, we inoculated replicated cyanogenic and acyanogenic genotypes with rhizobia. Cyanogenesis in white clover requires the production of two chemical precursors, cyanogenic glucosides and their hydrolyzing enzyme, linamarase. Cyanogenic genotypes produce both precursors (described as the AcLi cyanotype); acyanogenic genotypes lack one or both precursors (Acli, acLi, or acli cyanotypes). Here we only used plants with the acli cyanotype (lacking both precursors) to represent the acyanogenic morph, as they should have the lowest defense-related N-demand. We obtained genotypes from two biparental F_3_ mapping populations (DG and GS) that were segregating for cyanogenesis (Kuo et al. 2024; Kuo et al. 2026; Table S1). Using genotypes from these mapping populations allowed us to attribute any observed differences between cyanogenic and acyanogenic plants to cyanotype rather than artifacts of differences in their genetic backgrounds (particularly as cyanogenesis clines are frequently observed across environmental gradients {Kooyers & Olsen 2012; Kooyers et al. 2014; Santangelo et al. 2022}). We planted 20 stolon cuttings of each of 16 genotypes (8 genotypes per cyanotype) in pots (107 mL Cone-tainers™, Stuewe and Sons Inc., Corvallis, OR) that we disinfected and filled with a soil media consisting of two parts growing mix (Pro-Line HFEZ C/25 HydraFiber® Growing Mix, Jolly Gardener Products Inc., Poland, ME) and one-part calcine clay (DuraEdge® Fairball™, Grove City, PA). We disinfected all pots by soaking them in a broad range disinfectant (Physan 20™, Maril Products Inc., Tustin, CA) for 10 minutes and sterilized the soil media by running the soil mixture through a 30-minute sterilization cycle (minimum temperature: 121°C) in an autoclave three times with a 24 h rest period in between each cycle. We excluded cuttings that had not developed fully expanded leaves by the time of the first inoculation, so our experiment ultimately included 24 cyanogenic and 24 acyanogenic plants (48 total experimental plants) originating from 8 cyanogenic and 8 acyanogenic lines of clover. We grew the plants under a 16 h photoperiod at ∼24°C in a growth room at the University of Michigan Biological Sciences Building Plant Growth Facility. We watered plants daily for the duration of the experiment.

#### Rhizobium inoculation

We inoculated the plants with a strain *of Rhizobium leguminosarum* isolated from an old-field successional habitat at Kellogg Biological Station (Hickory Corners, MI, USA) (Strain #100 as isolated in Weese et al., 2015). To prepare the inoculum, we cultured rhizobia in liquid Tryptone Yeast media (5 g L^-1^ tryptone, 3 g L^-1^ yeast extract, 6.05 mM CaCl_2_ 2H_2_0) at 30°C for 48 hours. We diluted the inoculum with additional liquid media to a standard optical density (OD_620_) of 0.1 to standardize the number of rhizobium cells applied to each plant. We added 1 mL of diluted inoculum to each plant by pipetting inoculum onto the soil surface. To ensure rhizobium colonization, we inoculated each plant a second time three weeks after the initial inoculation using the same methods.

#### Data collection

We harvested plants 8 weeks after the second inoculation. To estimate differences in leaf tissue quality and N content, we measured leaf chlorophyll content using a SPAD 502 Plus Chlorophyll Meter (Spectrum® Technologies Inc., Aurora, IL). We measured leaf chlorophyll content on three haphazardly selected mature leaves of each plant and calculated an average leaf chlorophyll content per plant. We then harvested the above- and below-ground biomass of each plant. We dried the aboveground biomass of each plant at 60°C for at least 72 hours and then weighed them. We counted root nodules and estimated mean nodule mass by weighing 10 randomly selected nodules from each plant, or all nodules collected if a plant had less than 10 nodules. We estimated total nodule mass per plant by multiplying total nodule count by mean nodule mass. We then dried and weighed belowground biomass.

#### Data analysis

To test whether cyanogenesis affected plant performance or allocation to rhizobia, we used linear mixed effects models of above- and below-ground biomass, leaf chlorophyll content, mean nodule mass, total nodule mass, and total nodule count with main effects of cyanotype and mapping population (DG or GS) and a random effect of genotype using the “lmer” function in the *lme4* R package (Bates et al. 2015). To account for potential effects of plant size on investment in rhizobia, we also ran an additional model of nodule size that included aboveground plant biomass as a covariate. We log transformed all biomass and nodule data to meet the assumptions of normality. We performed all statistical analyses using R version 4.1.3 (R Core Team 2022).

### Experiment 2: Evolutionary responses of rhizobia to plant cyanotype

#### Experimental setup

We grew cyanogenic and acyanogenic clover in 500 mL clay pots (disinfected with Physan 20™, as in Experiment 1) filled with steam-pasteurized potting media (mixture of equal parts calcined clay (Turface®, Turface Athletics™, Buffalo Grove, IL, USA) and MetroMix820 (SunGro® Horticulture, Agawam, MA, USA) in greenhouses at Indiana University. To inoculate our initial mesocosms with a diverse soil community, we collected soil from old-field successional plots in the Main Cropping System Experiment (Treatment 7) at the Kellogg Biological Station Long-term Ecological Research Site (Hickory Corners, Michigan). We collected 20-cm deep cores of soil from six replicate old-field plots on January 10, 2022 and homogenized and sieved the soil to 4 mm. After storing the soil at 4°C for two days, we created a soil slurry by mixing water into the soil in an 8:3 ratio and inoculated each pot with 40 mL of the slurry. For all subsequent propagations, we created a new mesocosm by replacing 50% (v/v) of the conditioned soil from the previous propagation with fresh steam-pasteurized potting media and planting a new cutting of the same genotype.

To condition soils with plants of different cyanotypes, we used 12 cyanogenic and 12 acyanogenic lines of clover from the same two F3 mapping populations (DG and GS) that we used in our first experiment. Because clover is self-incompatible, we propagated all plants through *c.* 3 cm stolon cuttings in an autoclaved 1:1 mixture of vermiculite and perlite on a misting bench 15 days before planting into the mesocosms. We planted 8 replicate mesocosms per plant genotype and fertilized half of the replicates with urea at a rate of 11 g N/m^2^ applied in three equal aliquots at 2, 3, and 4 weeks after planting. To minimize contamination, we watered plants daily using a drip irrigation system. After 8 weeks of growth, we harvested the clover aboveground biomass and let the mesocosms sit for 21 days to allow the nodules to senesce and release *R. leguminosarum* into the soil. We then started the next iteration of clonal propagation by planting a new cutting in 50% (v/v) of soil from the mesocosm and repeated the conditioning process for a total of five 8-week plant growth cycles.

#### Trapping Rhizobia and assessing how cyanotype shapes the broader soil microbial community’s effect on plant growth

After harvesting the plants from the fifth cycle of plant conditioning, we allowed the nodules to senesce in the mesocosms for 21 days. We then prepared an inoculant by suspending 10 mL of the conditioned soil into 50 mL of deinoized water and pipetted 3 mL of this solution onto autoclaved potting media in a 107 mL conetainer. We planted *T. repens cv. Ladino* seeds (Ernst Seeds, Meadville, PA) to assess the effects of the conditioned soil microbial communities on plant growth and trap *R. leguminosarum*. We planted three surface-sterilized seeds (treated with 12% bleach) in each pot (thinning to one seedling after 21 d) and grew them in a growth room at the University of Michigan Biological Sciences Building Plant Growth Facility with a 16 h photoperiod. After growing the plants for 40 days, we dried and weighed above- and belowground biomass to assess the overall effects of microbial communities conditioned by the two plant cyanotypes on plant growth. Before drying the belowground biomass, we counted nodules and collected five randomly selected nodules from each plant to isolate *R. leguminosarum*. To culture *R. leguminosarum*, we surface-sterilized the nodules by dipping them briefly in 100% ethanol, soaking them in bleach for 90 s, and then streaking them on Tryptone-Yeast Extract media (Beringer 1974) with plates 1.8% Bacto-Agar (Becton, Dickinson and Company, Sparks, MD, USA) and incubating at 30°C. We isolated single colonies by serially replating and ultimately cultured 103 *R. leguminosarum* strains.

#### Assessing rhizobium quality

We assessed the quality of the evolved *R. leguminosarum* strains by inoculating them onto clover and assessing plant growth. We planted seeds of *T. repens cv. Ladino* in potting media (Pro-Line HFEZ C/25 HydraFiber® Growing Mix, Jolly Gardener Products Inc., Poland, ME) that had been autoclaved three times with a 24 h rest period between cycles. After 25 days, we cultured each strain in liquid TY media (Beringer 1974) and inoculated each strain onto three replicate plants with 1 mL of liquid culture (OD_620_ = 0.1). We inoculated plants a second time using the same methods one week later. We grew the plants under the same growing conditions as the rhizobium trapping plants and harvested the above- and below-ground biomass after 49-50 days. We counted the nodules on each plant and dried the biomass at 60°C for at least 72 hours before weighing.

#### Data analysis

To test if cyanotype affects rhizobium evolution, we ran linear mixed effects models with response variables of above- and below-ground biomass or nodule number with fixed effects of cyanotype (the cyanotype the rhizobium strain had evolved with) and mapping population (DG or GS) and random effects of strain and plant line (clover accession the rhizobium strain had evolved with) using the “lme4” and “lmerTest” R packages (Bates et al. 2015; Kuznetsova et al. 2017). Initially, we included whether the plants were fertilized or not during the conditioning phase as a fixed effect, but ultimately omitted this factor from models because it did not explain significant variation. We square root transformed nodule count data prior to statistical analyses to meet assumptions of normality. To further determine whether effects of cyanotype on nodule count were the result of differences in availability of carbon to invest between cyanotypes, we ran an additional model on nodule count with aboveground biomass as a covariate. To test if cyanotype affects plant growth through shaping the broader soil microbial community (rather than just individual rhizobium strains), we also ran linear mixed effects models with response variables of above- and below-ground biomass and nodule number of the plants that the rhizobium were trapped from, with fixed effects of conditioning cyanotype and population (DG or GS) and random effects of plant line and inoculant (conditioning mesocosm replicate). As six researchers harvested the plants, we also included harvester as a random effect in the model of nodule number to account for variation in nodule counting ability. We performed all statistical analyses using R version 4.1.3 (R Core Team 2022).

## Results

### Experiment 1: Effects of cyanotype on plant investment in rhizobia

Cyanogenic plants produced on average 79% larger nodules relative to acyanogenic plants (F_1,44_ = 5.78, p = 0.02; Table S2; Figure 1a), with no difference between populations (F_1,44_ = 0.04, p = 0.34; Table S2; Figure 1a). Estimated total nodule mass and nodule count did not differ between plant cyanotypes (total nodule mass: F_1,12_ = 3.03, p = 0.11; nodule count: F_1,11_ = 0.79, p = 0.39; Table S2; Figure 1b, c) or populations (both p > 0.35), although, cyanogenic plants tended to produce greater total nodule mass and more nodules than acyanogenic plants.

**Figure 1:**
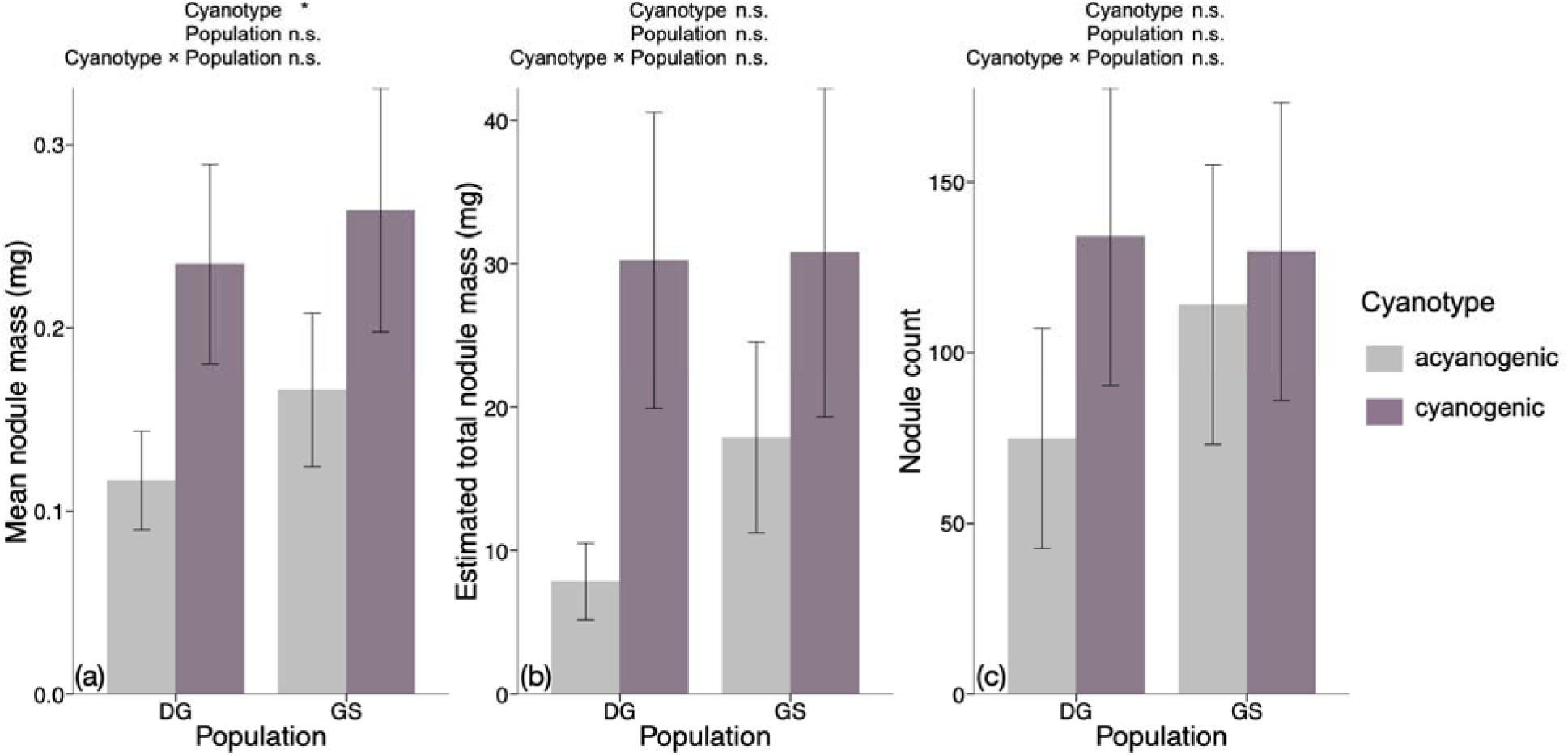
Mean nodule mass (mg; a), estimated total nodule mass (mg; b), and nodule count (c) of acyanogenic (gray bars) and cyanogenic (purple bars) clover from two biparental F_3_ mapping populations (“DG” and “GS”) inoculated with rhizobia. Values presented are estimated marginal means ± 1 SE. Significant effects of cyanotype are reported as: *p ≤ 0.05; **†**p < 0.1.

While cyanogenic plants were larger (Figure 2), the difference in nodule mass between cyanotypes remained significant even after accounting for variation in plant size (F_1,43_ = 6.06, p = 0.02; Table S3). Cyanogenic plants produced on average 60% more aboveground biomass than acyanogenic plants (F_1,11_ = 5.43, p = 0.039; Table S2; Figure 2a). Cyanogenic plants also produced on average 73% more belowground biomass than acyanogenic plants (F_1,10_ = 12.55, p = 0.006; Table S2; Figure 2b) with a marginal difference between populations (F_1,10_ = 3.65, p = 0.09; Table S2; Figure 2b). Leaf chlorophyll content (SPAD) did not differ between cyanotypes (F_1,12_ = 0.24, p = 0.63; Table S2; Figure 2c) with a marginal difference between populations (F_1,12_ = 4.43, p = 0.057; Table S2; Figure 2c).

**Figure 2:**
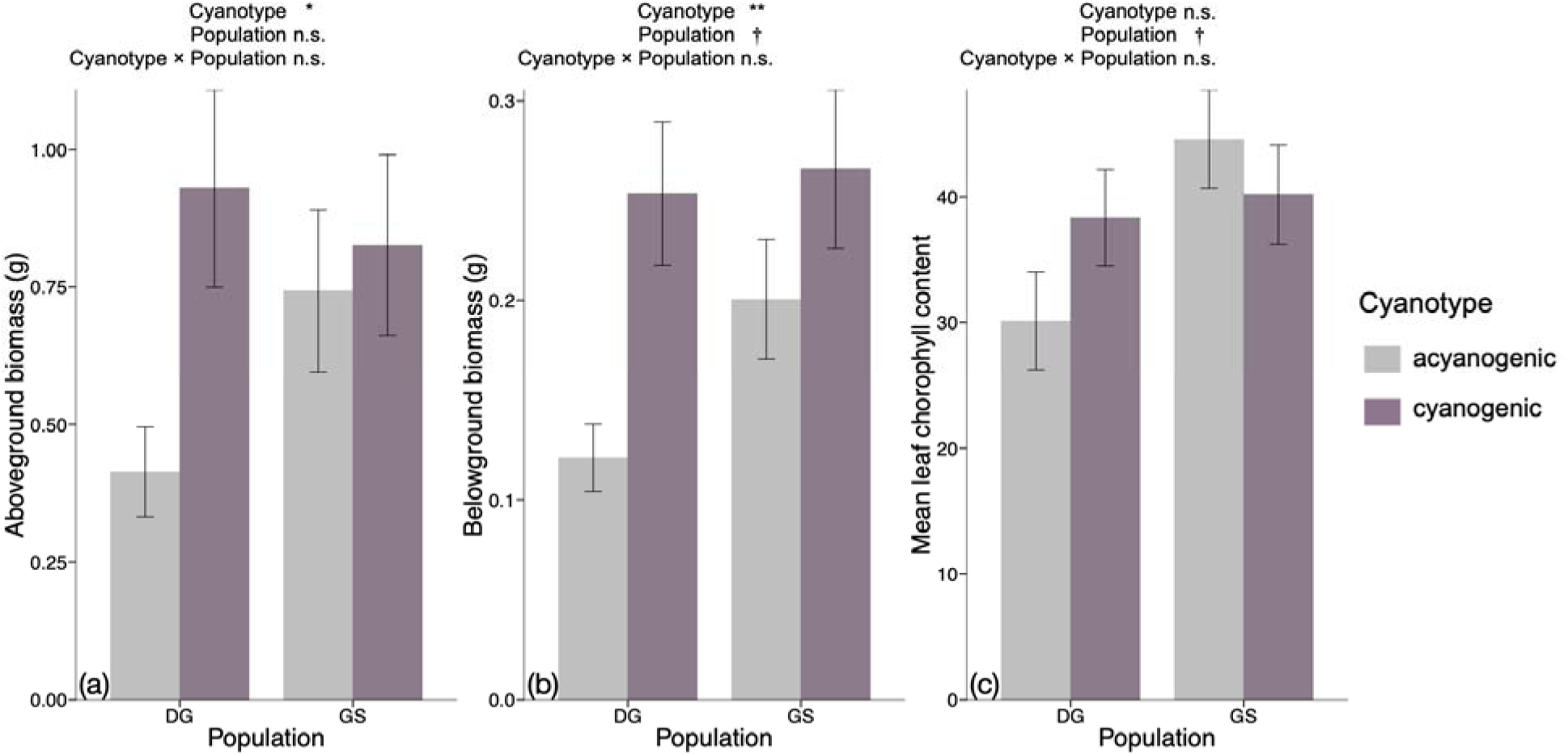
Aboveground biomass (g; a), belowground biomass (g; b), and mean SPAD (leaf chlorophyll content estimate; c) of acyanogenic (gray bars) and cyanogenic (purple bars) clover from two biparental F_3_ mapping populations (“DG” and “GS”) inoculated with rhizobia. Values presented are estimated marginal means ± 1 SE. Significant effects of cyanotype are reported as: **p ≤ 0.01; *p ≤ 0.05; **†**p < 0.1.

### Experiment 2: Effect of cyanotype on soil community quality and the evolution of rhizobium quality

#### Effect of cyanotype on soil microbial community effects on plant growth

We observed no difference in aboveground biomass produced by plants grown in whole soil microbial communities conditioned by the two cyanotypes (F_1,521_ = 0.01, p = 0.92; Table S4; Figure 3a). However, plants grown in soil communities conditioned by cyanogenic plants produced 11% more belowground biomass (F_1,518_ = 6.2, p = 0.01; Table S4; Figure 3b) and marginally more nodules (F_1,19_ = 3.63, p = 0.07; Table S4; Figure 3b) than those inoculated with soil communities conditioned by acyanogenic plants. The amount of aboveground biomass produced per nodule was similar between plants grown with soil communities conditioned by the two cyanotypes (F_1,19_ = 2.24, p = 0.15; Table S4; Figure 3d). Aboveground biomass, belowground biomass, nodule count, and the amount of aboveground biomass produced per nodule did not differ between populations (Table S4).

**Figure 3:**
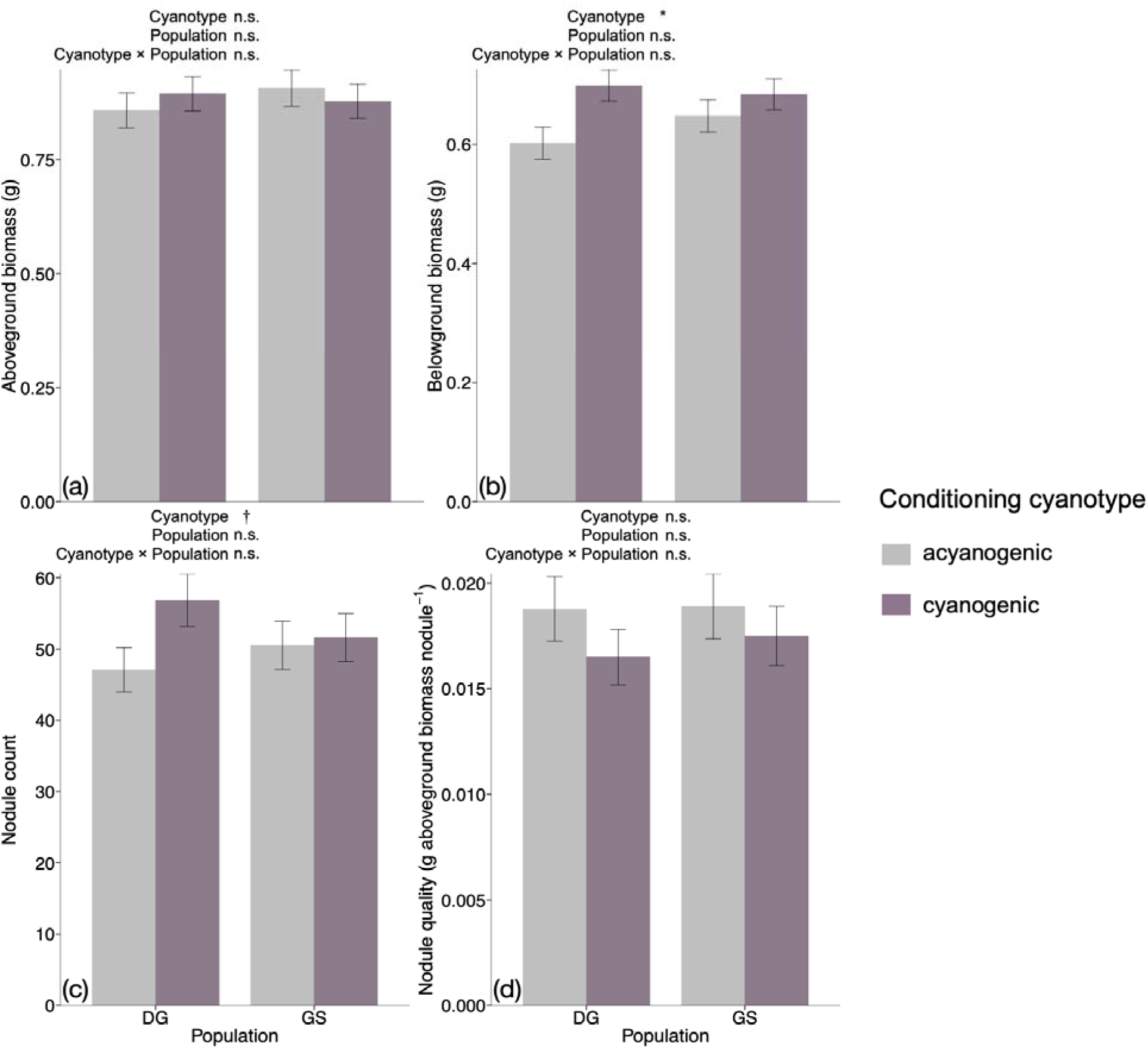
Aboveground biomass (g; a), belowground biomass (g; b), total nodule count (c), and nodule quality (g aboveground biomass nodule^-1^; d) of clover from two biparental F_3_ mapping populations (“DG” and “GS”) inoculated with whole soil microbial communities conditioned by acyanogenic (gray bars) or cyanogenic (purple bars) clover over five 8-week plant growth cycles. Values presented are estimated marginal means ± 1 SE.

#### Effect of cyanotype on the evolution of rhizobium quality

Aboveground biomass (F_1,98_ = 0.65, p = 0.42; Table S5; Figure 4a) and belowground biomass (F_1,99_ = 1.1, p = 0.3; Table S5; Figure 4b) did not differ between clovers inoculated with rhizobia that had evolved in association with cyanogenic clover lines and plants inoculated with rhizobia that had evolved in association with acyanogenic clover lines. Clover aboveground biomass (F_1,98_ = 0.03, p = 0.87; Table S5; Figure 4a) and belowground biomass (F_1,99_ = 0.54, p = 0.46; Table S5; Figure 4b) also did not differ between populations. The number of nodules produced on clovers was similar between rhizobia strains from soils conditioned by the two plant cyanotypes (F_1,18_ = 0.3, p = 0.59; Table S5; Figure 4c) and remained similar even after including aboveground biomass as a covariate in our analysis (F_1,18_ = 0.08, p = 0.78; Table S6; Figure 4c). The number of nodules produced on clovers did not differ between populations (F_1,18_ = 0.51, p = 0.48; Table S5; Figure 4c).

**Figure 4:**
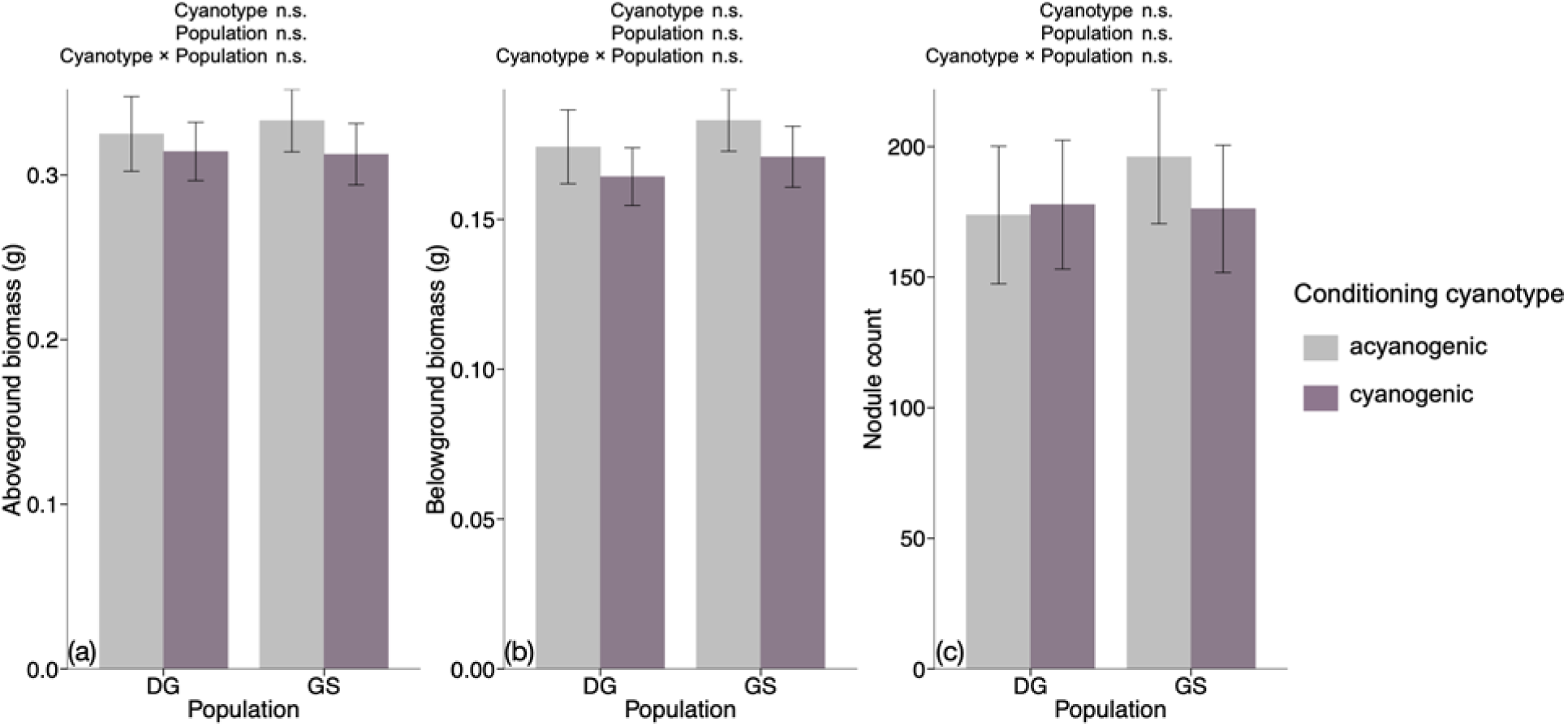
Aboveground biomass (g; a), belowground biomass (g; b), and total nodule count (c) of clover from two biparental F_3_ mapping populations (“DG” and “GS”) inoculated with rhizobium strains that had evolved in association with acyanogenic (gray bars) or cyanogenic (purple bars) clover for over five 8-week plant growth cycles. Values presented are estimated marginal means ± 1 SE.

## Discussion

### Cyanogenic plants invest more in resource mutualism

Here, we show that a plant defense trait can affect investment in microbial mutualists. Cyanogenic clover lines produced nodules that were more than twice the size of those of their acyanogenic counterparts. Importantly, our clover genotypes varied in the cyanogenesis trait but were otherwise from a homogeneous genetic background as F3 lines in pedigreed mapping populations descended from biparental crosses, suggesting that it is the production of cyanogenic compounds rather than other correlated traits causing the difference between cyanotypes. We also observed that cyanogenesis increased nodule size across both populations we tested, indicating the robustness of the effect. In addition, this greater nodule investment was not simply the result of cyanogenic plants being larger with more carbon to invest. While cyanogenic plants were larger, the difference in nodule size between cyanotypes remained significant even after accounting for variation in plant size. Greater N availability in the abiotic environment often corresponds with reduced investment in resource mutualists as plants can meet N demands without incurring the cost of supporting symbioses (i.e., Streeter & Wong 1988; Luciński et al. 2002; Glyan’ko et al. 2009; Friel & Friesen 2019). However, plant investment in resource mutualists can also vary with plant N demand. In lima bean (*Phaseolus lunatus*), highly cyanogenic plants had twice as many nodules as minimally cyanogenic plants (Godschalx et al. 2017), in line with our results showing that cyanogenic plants invest more in resource mutualists.

In addition to producing larger nodules, cyanogenic clover also accumulated greater above- and below-ground biomass relative to acyanogenic clover. Given how costly it is for plants to produce cyanogenic glucosides (Kaschuk et al. 2009), we were surprised that cyanogenic clovers were larger than acyanogenic clover. For example, Kempel et al. (2009) found that acyanogenic clovers were almost twice the size of their cyanogenic counterparts in the absence of rhizobia, indicating a substantial cost of cyanogenesis under N limitation. However, symbiosis with rhizobia enhanced the growth of the cyanogenic morphs enough to eliminate the size difference between the two cyanotypes (Kempel et al. 2009). Thus, forming symbioses with rhizobia can stimulate increased growth in cyanogenic plants despite the carbon cost of sustaining the mutualism (Kempel et al. 2009, Thamer et al. 2011). Here, we suggest that cyanogenic plants are investing more in rhizobia overall by producing larger nodules that support a greater density of mutualists, and associating with a greater number of symbionts may lead to greater biomass for cyanogenic plants.

Though cyanogenic clovers were larger, leaf greenness (a proxy for leaf chlorophyll content) did not vary between cyanogenic or acyanogenic lines. Thus, while increasing associations with rhizobia likely facilitated increased plant growth, rhizobium associations did not necessarily improve plant tissue quality. Here, cyanogenic plants seem to be investing the N acquired from their high density of mutualists into growth (and likely cyanogenesis), which would not be reflected in leaf greenness.

### Cyanogenesis does not affect the evolution of cooperative mutualists after 40 weeks of host genotype association

After evolving replicated rhizobium populations in association with cyanogenic and acyanogenic clover lines for 40 weeks (5 plant cycles), there was no difference in the quality of rhizobium mutualists. Cyanogenic plants also did not shape the broader soil microbial community in ways that affected plant growth differently than their acyanogenic counterparts. In this experiment, we originally inoculated cyanogenic and acyanogenic clover lines with field soil (homogenized from several old-field plots) that contained diverse microbial communities, including diverse rhizobium populations (Weese et al. 2015). Thus, there was likely substantial genetic variation in rhizobium quality for the clovers to select from. Other work has observed evolutionary shifts in rhizobium quality over several plant generations (Batstone et al. 2020; Doyle et al. 2026), though previous research has not tested the evolution of rhizobia quality explicitly in the context of plant defenses.

While we conducted our experiment in a greenhouse, legume-rhizobium mutualisms in the field exist within complex communities and ecosystems. Interactions between plants and their herbivores as well as light and other resource availability can modify the amount of available carbon a plant can allocate to resource mutualists (Lau et al. 2012; Simonsen & Stinchcombe 2014), and the degree to which plants are able to provide carbon to mutualists may in turn affect rhizobium evolution (Caple 2025). Additionally, plant community composition can shape plant investment in rhizobia (Zhou et al. 2022; Caple 2025). Here, we may not have observed an effect of cyanotype on rhizobium evolution because our experiment occurred in a simplified environment that eliminated the biotic and abiotic factors that likely would have further shaped clover-rhizobium interactions and rhizobium quality over time. The simplification of species interactions in our experiment is particularly relevant given that we were interested in how cyanotype, a plant phenotype that is explicitly related to and is a key mediator of plant-herbivore interactions, influences rhizobium evolution. For example, cyanogenic phenotypes not only have higher nitrogen demands but when herbivory is high, may also have more carbon to allocate to rhizobium symbionts compared to acyanogenic phenotypes that lose more leaf material to herbivores. This likely difference in herbivory would also be expected to promote the evolution of more cooperative mutualists in the presence of cyanogenic morphs.

While we did not find that cyanogenic plants’ greater investment in mutualism selected for higher quality rhizobia, their greater investment may increase the size of rhizobium populations in the soil. Repeated conditioning of the soil microbial community by the cyanogenic morph tended to increase nodulation on clover (Fig. 3c). Even though this increased nodulation did not correspond with improved plant growth in our experiment, perhaps such an increase in rhizobium investment could have a stronger impact in soils with lower titers of rhizobia such as farms where clovers have never been used as a cover crop. In agricultural systems, associations with beneficial microbes can promote plant growth, improve plant resistance to disease, and increase nutrient availability (Berg 2009). As such, planting cyanogenic morphs could be a strategy for increasing the abundance of rhizobia in such soils and enhancing cover crop productivity and biological N fixation.

Overall, our data show that cyanogenic clover lines invest more in N-fixing rhizobium mutualists, but that over 40 weeks of host association, they do not shape the evolution of more cooperative rhizobia relative to acyanogenic clover lines. While we show that cyanotype in part determines clover investment in rhizobia, future research should investigate whether N fertilization and cyanotype interactively affect plastic investment in rhizobia as well as the evolution of rhizobium quality overtime. Previous studies show that when legumes are fertilized with N, plant investment in rhizobia decreases (Luciński et al. 2002; Glyan’ko et al. 2009; Friel & Friesen 2019), but have not investigated whether this pattern depends on plant N demands or varies with cyanotype either in a single growing season or over longer time scales. This would be a particularly relevant area of future research given the vast amounts of N being input into agricultural systems annually (Ackerman et al. 2019). Improving our understanding of the relationship between plant traits like cyanogenesis, investment in resource mutualists, and the evolution of cooperation and quality of microbial mutualists could be important in helping us more effectively and sustainably manage widespread agricultural systems.

## Acknowledgments

This work was supported by a USDA NIFA grant (2022-67012-38587) and the University of Michigan Institute for Global Change Biology. We would like to thank Omilia Alicea, Caleb Callahan, Mackenzie Caple, Alexis Cooper, Madeleine Gellinger, Mark Hammond, Claire Kostecki, Madeleine Schouman, Avalon Yuhas, and Jordan Ziss for their help with measuring plants, counting nodules, and culturing rhizobia.

## References

Akçay, E., & Simms, E. L. (2011). Negotiation, sanctions, and context dependency in the legume-rhizobium mutualism. The American Naturalist, 178(1), 1–14.

Ackerman, D., Millet, D. B., & Chen, X. (2019). Global estimates of inorganic nitrogen deposition across four decades. Global Biogeochemical Cycles, 33(1), 100–107.

Bates, D., Mächler, M., Bolker, B., & Walker, S. (2015). Fitting linear mixed-effects models using lme4. Journal of statistical software, 67, 1–48.

Batstone, R. T., O’Brien, A. M., Harrison, T. L., & Frederickson, M. E. (2020). Experimental evolution makes microbes more cooperative with their local host genotype. Science, 370(6515), 476–478.

Berg, G. (2009). Plant–microbe interactions promoting plant growth and health: perspectives for controlled use of microorganisms in agriculture. Applied microbiology and biotechnology, 84(1), 11–18.

Beringer, J. E. (1974). R factor transfer in Rhizobium leguminosarum. Microbiology, 84(1), 188–198.

Bever, J. D. (2015). Preferential allocation, physio-evolutionary feedbacks, and the stability and environmental patterns of mutualism between plants and their root symbionts. New Phytologist, 205(4), 1503–1514.

Boter, M., & Diaz, I. (2023). Cyanogenesis, a plant defence strategy against herbivores. International Journal of Molecular Sciences, 24(8), 6982.

Caple, M. A. (2025). Cooperation Under Changing Conditions: Tests of Mutualism Theory in Legume-Rhizobium Systems (Doctoral dissertation, Indiana University).

Doyle, R. T., Su, X., Gallick, C., Blaszynski, M. M., Perry, E., Griesbaum, K., … & Heath, K. (2026). Negative frequency-dependent selection maintains partner quality variation in a keystone nutritional mutualism. bioRxiv, 2026–02.

Friel, C. A., & Friesen, M. L. (2019). Legumes modulate allocation to rhizobial nitrogen fixation in response to factorial light and nitrogen manipulation. Frontiers in Plant Science, 10, 1316.

Gleadow, R. M., & Møller, B. L. (2014). Cyanogenic glycosides: synthesis, physiology, and phenotypic plasticity. Annual review of plant biology, 65(1), 155–185.

Glyan’ko, A. K., Vasil’eva, G. G., Mitanova, N. B., & Ishchenko, A. A. (2009). The influence of mineral nitrogen on legume-rhizobium symbiosis. Biology Bulletin, 36(3), 250–258.

Godschalx, A. L., Tran, V., & Ballhorn, D. J. (2017). Host plant cyanotype determines degree of rhizobial symbiosis. Ecosphere, 8(9), e01929.

Hruska, A. J. (1988). Cyanogenic glucosides as defense compounds: a review of the evidence. Journal of Chemical Ecology, 14(12), 2213–2217.

Imakumbili, M. L., Semu, E., Semoka, J. M., Abass, A., & Mkamilo, G. (2019). Cyanogenic glucoside production in cassava: The comparable influences of varieties, soil moisture content and nutrient supply. bioRxiv, 649236.

Kaschuk, G., Kuyper, T. W., Leffelaar, P. A., Hungria, M., & Giller, K. E. (2009). Are the rates of photosynthesis stimulated by the carbon sink strength of rhizobial and arbuscular mycorrhizal symbioses?. Soil Biology and Biochemistry, 41(6), 1233–1244.

Kempel, A., Brandl, R., & Schädler, M. (2009). Symbiotic soil microorganisms as players in aboveground plant–herbivore interactions–the role of rhizobia. Oikos, 118(4), 634–640.

Kiers, E.T., Palmer, T. M., Ives, A. R., Bruno, J. F., & Bronstein, J. L. (2010). Mutualisms in a changing world: an evolutionary perspective. Ecology letters, 13(12), 1459–1474.

Kooyers, N. J., & Olsen, K. M. (2012). Rapid evolution of an adaptive cyanogenesis cline in introduced North American white clover (Trifolium repens L.). Molecular Ecology, 21(10), 2455–2468.

Kooyers, N. J., Gage, L. R., Al-Lozi, A., & Olsen, K. M. (2014). Aridity shapes cyanogenesis cline evolution in white clover (T Rifolium Repens L.). Molecular ecology, 23(5), 1053–1070.

Kuo, W. H., Cunningham, E., Guo, E., & Olsen, K. M. (2024). Genetics and plasticity of white leaf mark variegation in white clover (Trifolium repens L.). Annals of Botany, 134(6), 949–958.

Kuo, W. H., Small, L. L., & Olsen, K. M. (2026). Phenotypic and genetic bases of variable drought stress response in a widely adapted allotetraploid species. *American Journal of Botany*, e70200.

Kuznetsova, A., Brockhoff, P. B., & Christensen, R. H. (2017). lmerTest package: tests in linear mixed effects models. Journal of statistical software, 82, 1–26.

Lau, J. A., Bowling, E. J., Gentry, L. E., Glasser, P. A., Monarch, E. A., Olesen, W. M., … & Young, R. T. (2012). Direct and interactive effects of light and nutrients on the legume-rhizobia mutualism. Acta Oecologica, 39, 80–86.

Li, D., Wang, J., Chen, R., Chen, J., Zong, J., Li, L., … & Guo, H. (2024). Nitrogen acquisition, assimilation, and seasonal cycling in perennial grasses. Plant Science, 342, 112054.

Luciński, R., Polcyn, W., & Ratajczak, L. (2002). Nitrate reduction and nitrogen fixation in symbiotic association Rhizobium-legumes. Acta biochimica polonica, 49(2), 537–546.

Montesinos-Navarro, A. (2023). Nitrogen transfer between plant species with different temporal N-demand. Ecology Letters, 26(10), 1676–1686.

Pringle, E. G. (2016). Orienting the interaction compass: resource availability as a major driver of context dependence. PLoS biology, 14(10), e2000891.

R Core Team (2022). R: A language and environment for statistical computing. R Foundation for Statistical Computing, Vienna, Austria. https://www.R-project.org/.

Sachs, J. L., Mueller, U. G., Wilcox, T. P., & Bull, J. J. (2004). The evolution of cooperation. The Quarterly review of biology, 79(2), 135–160.

Santangelo, J. S., Ness, R. W., Cohan, B., Fitzpatrick, C. R., Innes, S. G., Koch, S., … & Lampei, C. (2022). Global urban environmental change drives adaptation in white clover. Science, 375(6586), 1275–1281.

Simonsen, A. K., & Stinchcombe, J. R. (2014). Herbivory eliminates fitness costs of mutualism exploiters. New Phytologist, 202(2), 651–661.

Simonsen, A. K., Han, S., Rekret, P., Rentschler, C. S., Heath, K. D., & Stinchcombe, J. R. (2015). Short-term fertilizer application alters phenotypic traits of symbiotic nitrogen fixing bacteria. PeerJ, 3, e1291.

Streeter, J., & Wong, P. P. (1988). Inhibition of legume nodule formation and N2 fixation by nitrate. Critical Reviews in Plant Sciences, 7(1), 1–23.

Thamer, S., Schädler, M., Bonte, D., & Ballhorn, D. J. (2011). Dual benefit from a belowground symbiosis: nitrogen fixing rhizobia promote growth and defense against a specialist herbivore in a cyanogenic plant. Plant and soil, 341(1), 209–219.

Weese, D. J., Heath, K. D., Dentinger, B. T., & Lau, J. A. (2015). Long-term nitrogen addition causes the evolution of less-cooperative mutualists. Evolution, 69(3), 631–642.

Wendlandt, C. E., Gano-Cohen, K. A., Stokes, P. J., Jonnala, B. N., Zomorrodian, A. J., Al-Moussawi, K., & Sachs, J. L. (2022). Wild legumes maintain beneficial soil rhizobia populations despite decades of nitrogen deposition. Oecologia, 198(2), 419–430.

Weyl, E. G., Frederickson, M. E., Yu, D. W., & Pierce, N. E. (2010). Economic contract theory tests models of mutualism. Proceedings of the National Academy of Sciences, 107(36), 15712–15716.

Zhou, J., Wilson, G. W., Cobb, A. B., Zhang, Y., Liu, L., Zhang, X., & Sun, F. (2022). Mycorrhizal and rhizobial interactions influence model grassland plant community structure and productivity. Mycorrhiza, 32(1), 15–32.

