## Supplemental Materials for "Cyanogenic clovers invest more in rhizobia but do not affect rhizobium evolution"

**Supplementary Material**

**Table S1:** *Trifolium repens* genotypes used in experiments. All genotypes are F3 lines derived from biparental crosses of cyanogenic and acyanogenic genotypes (Kuo et al., 2024; Kuo et al., 2026).

| Cyanotype | Genotype | F2 cross |
| --- | --- | --- |
| Cyanogenic | GS_F3_010 | GSSG004 |
|  | GS_F3_031 | GSSG017 |
|  | GS_F3_032 | GSSG019 |
|  | GS_F3_061 | GSSG036 |
|  | DG_F3_010 | DGGD006 |
|  | DG_F3_018 | DGGD008 |
|  | DG_F3_023 | DGGD010 |
|  | DG_F3_050 | DGGD021 |
| Acyanogenic | GS_F3_018 | GSSG008 |
|  | GS_F3_060 | GSSG035 |
|  | GS_F3_088 | GSSG056 |
|  | GS_F3_106 | GSSG064 |
|  | DG_F3_022 | DGGD009 |
|  | DG_F3_119 | DGGD051 |
|  | DG_F3_231 | DGGD173 |
|  | DG_F3_241 | DGGD191 |

**Table S2** Experiment 1: Summary of ANOVAs evaluating differences in aboveground biomass (g), belowground biomass (g), mean leaf chlorophyll content, mean single nodule mass (mg), estimated total nodule mass (mg), and nodule count between acyanogenic and cyanogenic clover cyanotypes inoculated with a single rhizobia strain from two biparental F_3_ mapping populations.

| Effect | Aboveground biomass (g) | Belowground biomass (g) | Leaf chlorophyll content | Single nodule mass (mg) | Total nodule mass (mg) | Nodule  count |
| --- | --- | --- | --- | --- | --- | --- |
| Fixed factors |  |  |  |  |  |  |
| Cyanotype | **5.43* (1,11)** | **12.55** (1,10)** | 0.24 (1, 12) | **5.78* (1, 44)** | 3.03 (1,12) | 0.79 (1, 11**)** |
| Population | 1.42 (1, 11) | **3.65† (1, 10)** | **4.43† (1, 12)** | 0.04 (1, 44) | 0.96 (1,11) | 0.13 (1,11) |
| Random Effects |  |  |  |  |  |  |
| Genotype | 1.63 | 0 | 1.76 | 0 | 0.13 | 1.77 |

Table entries are F values (Numdf, Dendf). “Cyanotype” is cyanotype (acyanogenic or cyanogenic) of experimental plant. “Population” is the biparental F_3_ mapping population from which an experimental plant originates (DG or GS). “Genotype” is genotype of experimental plant. Significance of random effects was determined with likelihood ratio tests. Symbols indicate levels of statistical significance: **p ≤ 0.01; *p ≤ 0.05; **†**p < 0.1. Bold values indicate results with P-values < 0.1.

**Table S3** Experiment 1: Summary of ANOVAs evaluating differences in mean single nodule mass (mg) and estimated total nodule mass (mg) between acyanogenic and cyanogenic clovers inoculated with a single rhizobia strain from two biparental F_3_ mapping populations, while accounting for variation in plant size.

| Effect | Single nodule mass (mg) | Total nodule mass (mg) |
| --- | --- | --- |
| Fixed factors |  |  |
| Cyanotype | **6.06* (1, 43)** | 3.03 (1,12) |
| Population | 1.18 (1, 43) | 0.96 (1, 11) |
| AG biomass | 0.39 (1, 43) | 0.94 (1, 43) |
| Random Effects |  |  |
| Genotype | 0 | 0.11 |

Table entries are F values (Numdf, Dendf). “Cyanotype” is cyanotype (acyanogenic or cyanogenic) of experimental plant. “Population” is the biparental F_3_ mapping population from which an experimental plant originates (DG or GS). “AG biomass” is the aboveground biomass of the experimental plant. “Genotype” is genotype of experimental plant. Significance of random effects was determined with likelihood ratio tests. Symbols indicate levels of statistical significance: *p ≤ 0.05. Bold values indicate results with P-values < 0.1.

**Table S4** Experiment 2 (Trapping Generation): Summary of ANOVAs evaluating differences in aboveground biomass (g), belowground biomass (g), and nodule count, and nodule quality (g aboveground (“AG”) biomass nodule^-1^) of clover from two biparental F_3_ mapping populations that the rhizobia were trapped from to use in the final generation of Experiment 2.

| Effect | Aboveground biomass (g) | Belowground biomass (g) | Nodule  count | Nodule quality (g AG biomass nodule^-1^) |
| --- | --- | --- | --- | --- |
| Fixed factors |  |  |  |  |
| Cyanotype | 0.01 (1, 521) | **6.20* (1, 518)** | **3.63† (1, 19)** | 2.24 (1, 19) |
| Population | 0.18 (1, 521) | 0.33 (1, 518) | 0.06 (1, 20) | 0.22 (1, 19) |
| Harvester | n/a | n/a | **11.79*** (6, 495)** | **20.46*** (6, 508)** |
| Random Effects |  |  |  |  |
| Genotype | 0 | 0 | 0.01 | 0.49 |
| Inoculant | 0 | 0 | 0.06 | 0 |

Table entries are F values (Numdf, Dendf). “Cyanotype” is cyanotype (acyanogenic or cyanogenic) of experimental plant. Population” is the biparental F_3_ mapping population from which an experimental plant originates (DG or GS). “Harvester” is the identity of one of the 6 individuals that harvested experimental plants for this experiment. “Genotype” is genotype of the plant that conditioned the soil. “Inoculant” is which pot the rhizobia inoculant originated from. Symbols indicate levels of statistical significance: ***p ≤ 0.001; *p ≤ 0.05; **†**p < 0.1. Bold values indicate results with P-values < 0.1.

**Table S5** Experiment 2: Summary of ANOVAs evaluating differences in aboveground biomass (g), belowground biomass (g), and nodule count of clover inoculated with soil microbial communities conditioned by acyanogenic or cyanogenic clover from two biparental F_3_ mapping populations for five 8-week plant growth cycles.

| Effect | Aboveground biomass (g) | Belowground biomass (g) | Nodule  count |
| --- | --- | --- | --- |
| Fixed factors |  |  |  |
| Cyanotype | 0.65 (1, 98) | 1.10 (1, 99) | 0.30 (1, 18) |
| Population | 0.03 (1, 98) | 0.54 (1, 99) | 0.51 (1, 18) |
| Harvester | n/a | n/a | **14.83*** (5, 266)** |
| Random Effects |  |  |  |
| Genotype | 0 | 0 | 0 |
| Strain | **4.15*** | **4.51*** | **21.96***** |

Table entries are F values (Numdf, Dendf). “Cyanotype” is cyanotype (acyanogenic or cyanogenic) of experimental plant. Population” is the biparental F_3_ mapping population from which an experimental plant originates (DG or GS). “Harvester” is the identity of one of the 6 individuals that harvested experimental plants for this experiment. “Genotype” is genotype of the plant that the strain evolved with/was trapped from. “Strain” is the isolated rhizobia that a plant was inoculated with after five 8-week plant growth cycles. Symbols indicate levels of statistical significance: ***p ≤ 0.001; *p ≤ 0.05. Bold values indicate results with P-values < 0.1.

**Table S6** Experiment 2: Summary of ANOVA evaluating differences in nodule count of clover with soil microbial communities conditioned by acyanogenic or cyanogenic clover two biparental F_3_ mapping populations for five 8-week plant growth cycles while accounting for variation in plant size.

| Effect | Nodule  count |
| --- | --- |
| Fixed factors |  |
| Cyanotype | 0.08 (1, 18) |
| Population | 0.57 (1, 18) |
| Harvester | **21.12*** (5, 269)** |
| AG biomass | **38.60*** (1, 289)** |
| Random Effects |  |
| Genotype | 0 |
| Strain | **18.51***** |

Table entries are F values (Numdf, Dendf). “Cyanotype” is cyanotype (acyanogenic or cyanogenic) of experimental plant. Population” is the biparental F_3_ mapping population from which an experimental plant originates (DG or GS). “Harvester” is the identity of one of the 6 individuals that harvested experimental plants for this experiment. “AG biomass” is the aboveground biomass of the experimental plant. “Genotype” is genotype of the plant that the strain evolved with/was trapped from. “Strain” is the isolated rhizobia that a plant was inoculated with after five 8-week plant growth cycles. Symbols indicate levels of statistical significance: ***p ≤ 0.001. Bold values indicate results with P-values < 0.1.
